# Exploring the human cerebral cortex using confocal microscopy

**DOI:** 10.1101/2021.07.16.452651

**Authors:** Luca Pesce, Annunziatina Laurino, Marina Scardigli, Jiarui Yang, David A. Boas, Patrick R. Hof, Christophe Destrieux, Irene Costantini, Francesco Saverio Pavone

**Author notes:** Luca Pesce and Annunziatina Laurino contributed equally to this work.

## Abstract

Cover-all mapping of the distribution of neurons in the human brain would have a significant impact on the deep understanding of brain function. Therefore, complete knowledge of the structural organization of different human brain regions at the cellular level would allow understanding their role in the functions of specific neural networks. Recent advances in tissue clearing techniques have allowed important advances towards this goal. These methods use specific chemicals capable of dissolving lipids, making the tissue completely transparent by homogenizing the refractive index. However, labeling and clearing human brain samples is still challenging. Here, we present an approach to perform the cellular mapping of the human cerebral cortex coupling immunostaining with SWITCH/TDE clearing and confocal microscopy. A specific evaluation of the contributions of the autofluorescence signals generated from the tissue fixation is provided as well as an assessment of lipofuscin pigments interference. Our evaluation demonstrates the possibility of obtaining an efficient clearing and labeling process of parts of adult human brain slices, making it an excellent method for morphological classification and antibody validation of neuronal and non-neuronal markers.

## 1. Introduction

The characterization of both normal and abnormal expression of specific proteins and metabolic products using immunohistochemistry has allowed to explore the maturation and define the neuronal, glial, and blood vessels organization of the nervous system. However, processing whole brain regions into thin slices limits the investigation of such samples, introducing artifacts and affecting the detection of specific biomolecules (Silvestri et al., 2016). Recent advances in clearing methods for the purpose of histological analysis are becoming one of the main tools used for biological investigation of several kinds of tissues and organs. The common idea behind these approaches is to preserve proteins and nucleic acids, remove lipids, and render the specimens completely transparent by homogenizing the refractive index (Costantini et al., 2019; Tainaka et al., 2016). Different optical clearing methods have been developed to achieve high structural preservation, efficient probe penetration, and reduce light scattering. They can be categorized into three distinct groups: hydrophilic, hydrophobic, and tissue transformation or hydrogel-based methods (Silvestri et al., 2016; Ueda et al., 2020). Although dissolving cellular membrane lipids provides high tissue transparency, specific solvents and detergents used in hydrophilic and hydrophobic methods can induce protein loss and a high degree of denaturation. To overcome these drawbacks, tissue transformation approaches stabilize proteins and nucleic acids by cross-linking them to a hydrogel (Ueda et al., 2020). Methods like CLARITY (Chung et al., 2013), SWITCH (Murray et al., 2015), SHIELD (Park et al., 2019), and expansion microscopy (Chen et al., 2015) allow clearing and labeling any kind of tissues and organs. However, obtaining high-resolution information of the post-mortem human brain is still challenging owing to a number of factors such as post-mortem interval variability, intrinsic tissue properties, and the impossibility to use fluorescent proteins. Also, unblocked free aldehyde groups introduced during the fixation step may lead to high background autofluorescence throughout the tissue. Besides, the human brain is rich in blood and lipopigments (Ueda et al., 2020) – like lipofuscin (LF) – which can be used as natural landmarks (Costantini et al., 2021), but may interfere with the immunostaining process, generating non-specific signal. Here, we propose to define the molecular profile and morphology of neurons in processed human cortex slices using confocal microscopy in combination with the SWITCH/TDE clearing procedure (Costantini and Mazzamuto et al., 2021). The structural preservation of the SWITCH protocol (Murray et al., 2015) combined with the clarifying agent 2’2-thiodiethanol (TDE) (Costantini et al., 2015), enables discrimination of neuronal markers, glial cells, and microvasculature for post-mortem human brain characterization. Here, we characterize (1) the tissue autofluorescence in normal and transformed slices, (2) LF clustering, and (3) we probe several antibodies to assess their compatibility with SWITCH/TDE-processed slices using confocal microscopy.

## 2. Methods

### 2.1 Human brain specimens

Healthy human tissue samples were obtained from body donation programs (Association des dons du corps) of the Université de Tours and from the Massachusetts General Hospital (MGH). Prior to death, participants gave written consent for using their entire body, including the brain, for any educational or research purpose in which an anatomy laboratory is involved. The authorization documents of the Association des dons du corps was kept in the files of the Body Donation Program, while the informed consent authorization documents of MGH are kept in the files of MGH Autopsy Services in Boston, MA, and are available upon request. The tissue samples provided through MGH Autopsy Services do not require specific ethics approval documentation because they are collected within the general frame of the approved IRB submission to the Partners Institutional Biosafety Committee (PIBC, Protocol 2003P001937).

Sample 1 was injected in both carotid arteries with 4% buffered formalin the day after death. The brain was extracted the day after an immersed in the same fixative solution at RT and store for 6 months. Sample 2 was removed from the skull after death and placed in 10% neutral buffered formalin (pH 7.2-7.4; Diapath, Martinengo, Italy) for 3 months and then store at room temperature (RT) for 7 years in Periodate-lysine-paraformaldehyde 2% (PLP). Blocks from the fixed samples were washed in a phosphate-buffered saline (PBS) solution at 4 °C with gentle shaking for one month. Blocks were embedded in low melting agarose (4% in 0.01 M PBS) and cut into 450±50 μm-thick coronal sections with a vibratome (Vibratome 1000 Plus, Intracel, UK). After cutting, the agarose surrounding each slice was removed. For this work, we used two specimens: the precentral cortex of a 99-year-old (female) (sample 1), 6 months in formalin, was used for autofluorescence analysis, LF characterization, and immunolabeling validation; the superior frontal cortex of a 68-year-old (male), 7 years in formalin (sample 2) was used for immunolabeling validation. Both cases were neurotypical and had no neuropathological diagnosis of neurodegenerative disorders.

### 2.2 SWITCH protocol for human brain slices and TDE refractive index matching

The SWITCH protocol from Costantini and Mazzamuto et al., (2021) was used to clear large portions of the human brain samples. Briefly, 500 μm-thick slices were incubated for 24 h in a Switch-Off solution containing 50% PBS titrated to pH 3 using HCl, 25% 0.1 M HCl, 25% 0.1 M potassium hydrogen phthalate (KHP), and 4% glutaraldehyde at 4 °C. In this step, the protonation process of the primary amine groups prevents the reaction between glutaraldehyde and endogenous biomolecules. Next, the cross-linking reaction was performed by replacing the Switch-Off solution with PBS pH 7.4 with 1% glutaraldehyde 4 °C for 24 h. After 3 washes in 0.01 M PBS + 0.1% Triton (PBST) at room temperature (RT), the transformed tissue was inactivated by incubation overnight in a solution containing 4% glycine and 4% acetamide at 37 °C. Following 3 washes in PBST at RT, the slices were incubated in the clearing solution consisting of 200 mM SDS, 10 mM lithium hydroxide, 40 mM boric acid, for 7 days at 53 °C. The clearing solution was exchanged every 2 days. Following the clearing step, the transformed samples were extensively washed in PBST at 37 °C for 24 h. SWITCH-processed slices were finally cleared in a solution of increasing concentration of TDE, 20, 40, and 68% in PBS, 1 h in each incubation at RT with gentle shaking.

### 2.3 Tissue autofluorescence characterization

Cleared samples were imaged at 405 nm, 488 nm, and 561 nm using a Nikon C2 laser scanning confocal microscope with a Plan Fluor 60X 1.49 NA oil immersion objective. To quantify the fluorescence intensities in each channel, 4 regions of interest (ROI) of 10 x 10 μm were selected randomly in the grey matter using Fiji. Laser power at 405, 488, and 561 nm was set at 1 mW. Means and standard deviations were plotted using Origin. Statistical significance was assessed with two-way ANOVA followed by the Bonferroni post hoc test.

### 2.4 Lipofuscin characterization

For LF size characterization, SWITCH-processed samples were acquired using a Nikon C2 laser scanning confocal microscope with a Plan Fluor 60X 1.49 NA oil immersion objective at 405 nm, 488 nm, and 561 nm maintaining the same laser power (1 mW). Next, the area of LF granules was calculated using Fiji by manual segmentation of 40 distinguishable single clusters. After the selection of the LF clusters, their mean intensity, and standard deviation were calculated for each channel using Fiji.

To estimate LF area, Fiji was used to segment manually 20 neurons labeled with NeuN and 20 LF aggregates in the cell soma. Descriptive statistics, including mean, median, min-max, and 90th percentile were calculated using Origin. The images were acquired using a Nikon C2 laser scanning confocal microscope with a Plan Fluor 60X 1.49 NA oil immersion objective at 488 and 561 nm. For the intensity profile, Fiji was used to calculate the signal along with the cell soma in NeuN-immunostained and unstained samples, both marked with DAPI. For the signal-to-noise analysis, ROIs of 2 x 2 μm were selected in the cellular soma (LF and NeuN staining, N =30) and background (N=30) in Fiji.

### 2.5 Immunofluorescence labeling

For the labeling process, primary antibodies (SST, Abcam, ab30788; PV, Abcam, ab11427; VIP, Abcam, ab214244; NPY, Abcam, ab112473; CR, Proteintech, 12278-1-AP; GAD67, Abcam, ab26116; MAP2, Abcam, ab5392; SMI32, Merck, NE1023; NeuN, Merck, ABN91; β-Tub, Abcam, ab18207; Iba1, Abcam, ab195031; GFAP, Abcam, ab194324; Vim, Abcam, ab8069; Coll IV, Abcam, ab6586; GluS, Merck, MAB302; p-S6RP, Cell signaling, 2215; Lam B1, Abcam, ab16048) are incubated in the PBST solution for three days at 4 °C with a dilution of 1:50 for NeuN and 1:200 for the rest primary antibodies. After three washing steps in the PBST solution, each for 10 min, secondary antibodies (goat anti-chicken IgY AF 568, Abcam, ab175711; goat anti-chicken IgY AF 488, Abcam, ab150169; donkey anti-mouse IgG AF 568, Abcam, ab175700; donkey anti-rabbit IgG AF 568, Abcam, ab175470; goat anti-rabbit IgG AF 488, Abcam, ab150077; donkey anti-sheep IgG AF 568, Abcam, ab175712) with a dilution of 1:200 were incubated in PBST for two days at RT. After three washing steps in PBST at RT for 10 min each, the stained samples were equilibrated in the TDE/PBS optical clearing solution. Before imaging, DAPI was supplemented in the 68% TDE/PBS solution at the dilution 1:100 for 1 h. The images were acquired using a Nikon C2 laser scanning confocal microscope with a XLUMPlanFI 20X 0.95 NA water immersion objective at 405, 488, and 561 nm excitation. The maximum intensity projection (MIP) was performed using Fiji.

## 3. Results

### 3.1. SWITCH-processed slices show high autofluorescence signal

For probing the molecular composition of several neuronal classes in large samples of human brain specimens, the tissue needs to preserve the endogenous biomolecules and their antigenicity. To achieve this goal, we used a modified version of the SWITCH (system-wide control of interaction time and kinetics of chemicals; Murray et al., 2015) method that allows stabilization of the tissue architecture by using a set of buffers. The SWITCH-Off buffer (pH 3) suppresses the formation of the intra- and intermolecular cross-linking mediated by glutaraldehyde, uniformly to disperse the fixative molecules into the tissue. Then, the sample is moved to SWITCH-On buffer (pH 7) to initialize a homogenous tissue fixation. Then, the SWITCH-processed specimens are delipidated using a high temperature and permeabilized using sodium dodecyl sulfate (SDS) and Triton X-100 (Fig. 1A). The main advantage of this protocol is its compatibility with human specimens as glutaraldehyde generates a uniform fixation without the need of a perfusion step. However, the SWITCH-processed sample presents a drawback: the autofluorescence signal is increased. We characterized the modification of the autofluorescence signal after the procedure and observed that it was particularly enhanced at 488 and 561 nm by 3 and almost 4 times, respectively, with respect to the post-mortem fixed sample (Fig. 1B, ***P<0.001 vs non-processed tissue/TDE). These findings are in line with the literature (Lee et al., 2013), confirming that glutaraldehyde produces a red-shifted emission upon binding to peptides and proteins, in particular lysine residues, making it challenging to label specific neuronal markers in the human brain.

**Fig. 1.**
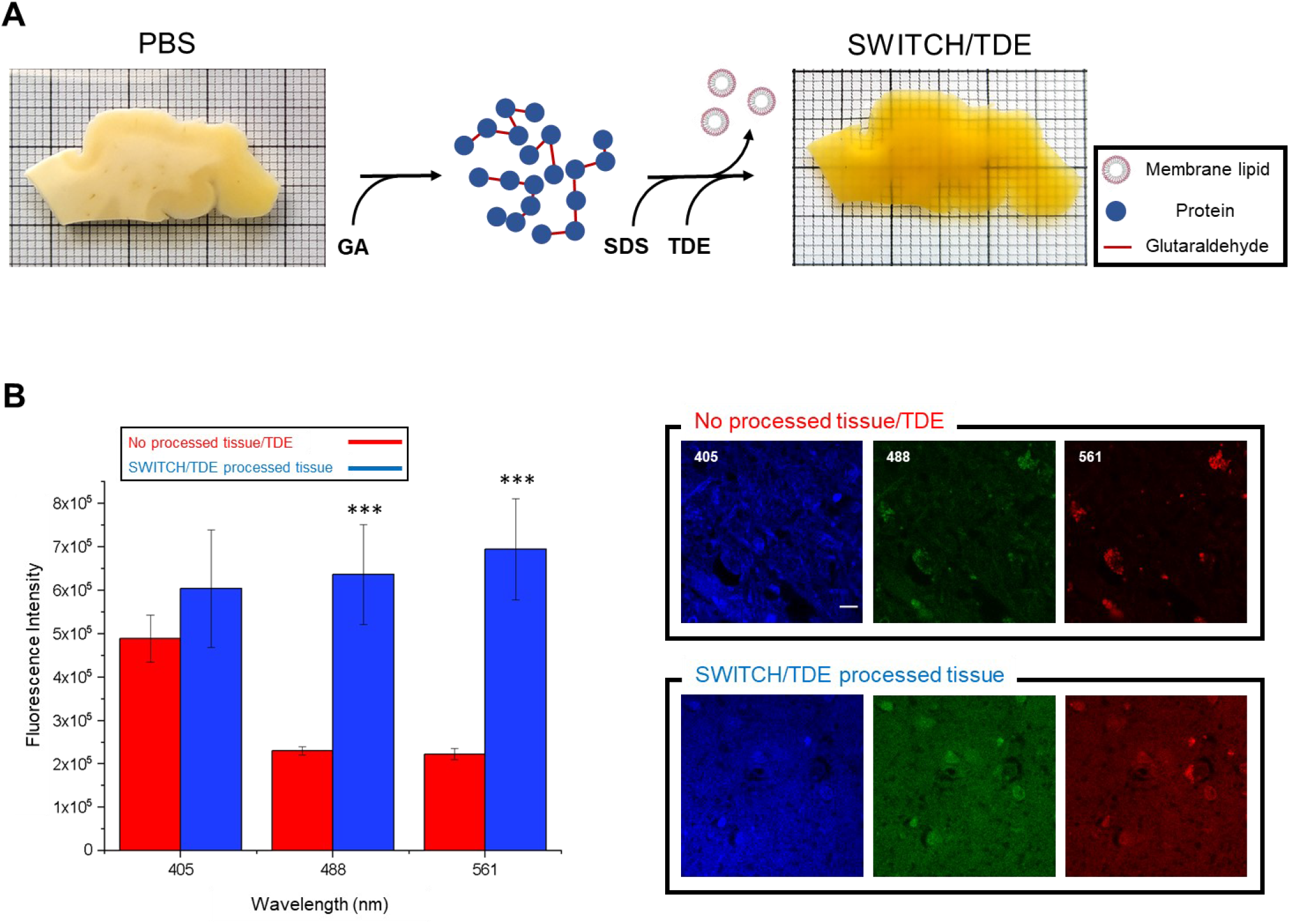
Schematic illustration of the clearing process. (A) 500 μm-thick slice of an adult human brain sample (99-years old; 6 months in formalin; precentral cortex) before and after the tissue treatment. First, the glutaraldehyde molecules diffuse into the sample, generating a tissue-gel hybrid. Then, membrane lipids are removed using high temperature and SDS. Finally, the sample is incubated in the TDE/PBS solution for refractive index matching. (B) Autofluorescence comparison between SWITCH/TDE and normal/TDE samples (N = 4). from various excitation wavelengths (405, 488, 561 nm). Laser power 1 mW. ***P<0.001 vs no processed tissue/TDE. Scale bar 10 μm.

### 3.2. Lipofuscin autofluorescence

The presence of other entities could interfere with the immunolabeling process and generate false positives. The human brain is rich in blood and autofluorescent lipopigments such as LF and neuromelanin (Monici, 2005), which are difficult to clear or decolorize. Compared to neuromelanin, which generates fluorescence upon exposure to UV and in the presence of peroxides or periodic acid (Elleder and Borovanský, 2001), LF is a strong autofluorescence emitter. As the accumulation of LF increases during aging (Terman and Brunk, 1998), we characterized this pigment in the brain of a very old patient (99 years old; Fig. 1A). To characterize the impact of LF fluorescence emissions, we estimated the amount of LF pigments. As shown in Figure 2A, we found that the human cortex contains a high concentration of LF, which presents as granule-shaped intracellular clusters ranging in size from 0.7 to 2.1 μm, with a red-shifted emission spectrum (Fig. 2A and 2B). Using light microscopy combined with immunolabeling, this endogenous autofluorescence could invalidate the proper recognition of labeled neurons, causing an incorrect interpretation of such images. As the laser-scanning microscope revealed, LF occupied part of the cell soma with respect to the more homogenous labeling obtained by NeuN immunostaining (see line profile, Fig. 2D). To avoid misinterpretation and to quantify the volume occupied by LF in the soma more accurately, it is possible to exploit the characteristic of LF that shows a higher fluorescence emission at 561 nm with respect to 488 nm. Indeed, to quantify the volume occupied by LF in the soma, neurons can be labeled with NeuN using the Alexa Fluor 488 to separate it spectrally from the LF signal. Using this approach it is possible to discriminate both contributions: LF shows a prominent localization in the proximity of the nucleus, distributing its signal in an area of about 30% of the cellular soma, while NeuN fully stains the neuronal body (Fig. 2E, in green). Although the signal-to-noise ratio of LF and NeuN immunostaining does not show a significant difference at 561 nm (Fig. 2C), the antibody staining highlights a complete distribution of the signal throughout the cytoplasm. With respect to the NeuN immunostaining, the differences between LF distribution and the labeled interneurons and glial cells are more evident. Such optimization allows the identification of the dendritic arbors of different cell types (e.g., parvalbumin-expressing neurons and glial cells) compared to the inhomogeneous and variable LF distribution (see the maximum intensity projection, Fig. 2F). Using these precautions, the LF-derived signal can be efficiently distinguished in the stained neurons and interneurons, avoiding false positives during image analysis.

**Fig. 2.**
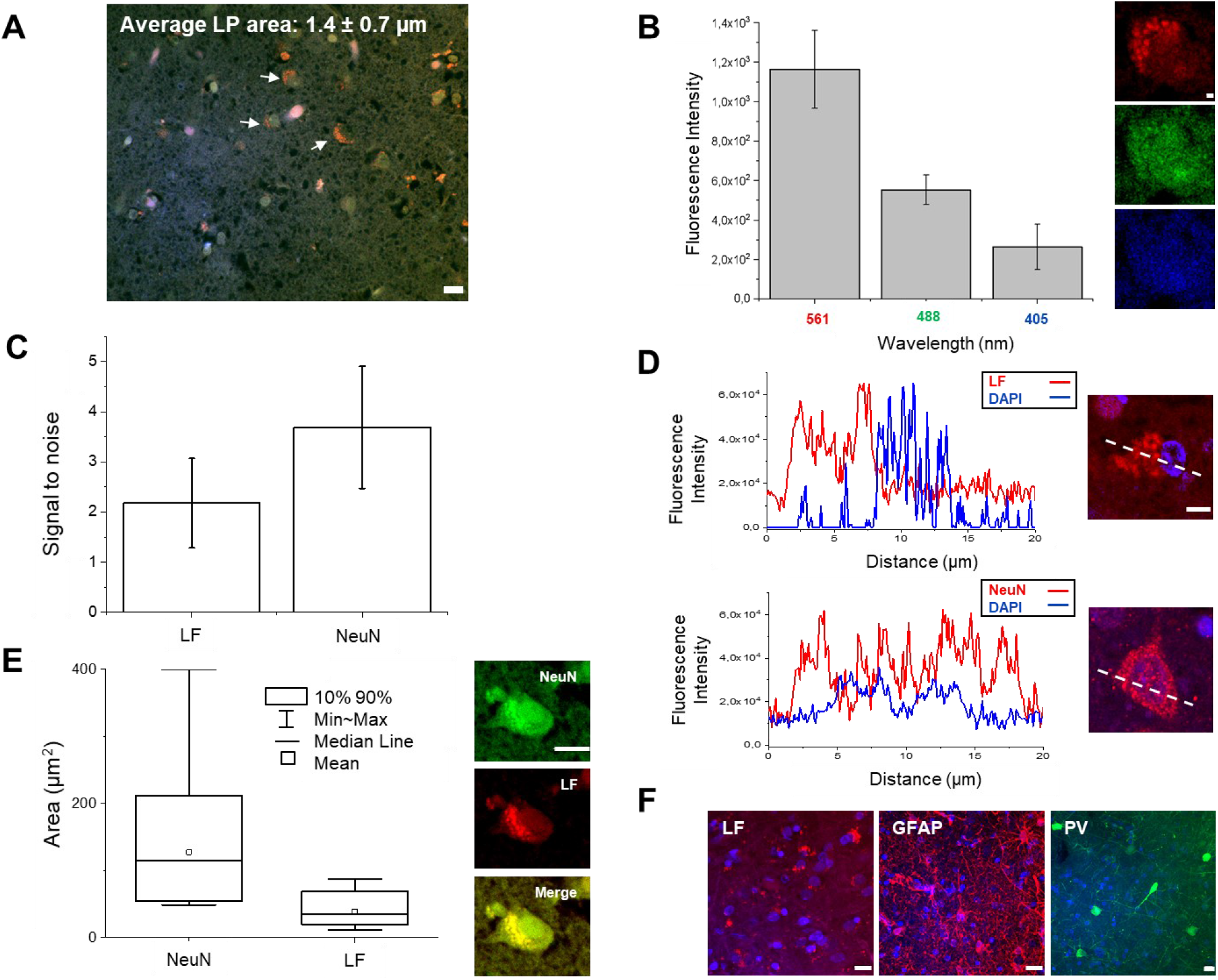
Lipofuscin characterization. (A) Average size of LF granular shape (pointed with white arrows) resolvable using confocal microscopy. Scale bar = 10 μm. (B) Plot of LFs emission at various excitation light (405, 488, 561 nm); excitation light 1 mW. Representative images of a neuron imaged at the three different wavelengths (from the top: 561, 488, 405nm). Scale bar = 1 μm. (C) Signal-to-noise ratio between the unstained (LF signal) and stained sample using NeuN antibody marked with Alexa Fluor 568. Both samples are labeled with DAPI. Excitation power of 1 mW at 561 nm. (D) Plot profile comparison between LF and NeuN/Alexa Fluor 568 labeled samples, both samples marked with DAPI. Scale bar = 5 μm. (E) Mean, median, min-max, and 90th percentile of 20 Neurons labeled for NeuN with Alexa Fluor 488 and LF aggregates. Scale bar = 10 μm. (F) Maximum intensity projection (MIP) of unstained sample (LF), which shows the LFs distribution, and two specimens labelled for GFAP and parvalbumin (PV) with Alexa Fluor 568 and Alexa Fluor 488, respectively. Scale bar = 10 μm.

### 3.3 Morphological classification using confocal microscopy

After fluorescence and distribution characterization of LF, we probed several antibodies using confocal microscopy for molecular cell-type characterization in the human cortex. To do this, we performed the immunostaining process on two elderly human brain specimens (99 and 68 years old, see Fig. 1A and Fig. 3A) kept in formalin for different periods of time before tissue processing (6 months and 7 years, respectively). To classify neocortical interneurons - neuronal subclasses which are less abundant than pyramidal cells in the neocortex - we used three different markers (Tremblay et al., 2016; DeFelipe et al., 2013). In particular, we selected non-overlapping populations of neocortical interneurons, which are characterized by the expression of the calcium-binding protein parvalbumin (PV), the neuropeptide somatostatin (Sst), and ionotropic serotonin receptor 5HT3a (5HT3aR). In addition, 40% of 5HT3aR-expressing interneurons also express vasointestinal peptide (VIP), which is not expressed in PV- or Sst-immunoreactive neurons (Tremblay et al., 2016; Zeng and Sanes, 2017). Based on these considerations, we first evaluated the compatibility of our clearing method for detecting these neuronal markers in processed human brain slices. Images of immunolabeled interneurons for Sst, PV, and VIP were acquired using a confocal microscope (Fig. 3B). In agreement with other works, PV-expressing cells show more complex dendritic and axonal arborization patterns than Sst-expressing neurons in the human neocortex. Next, other antibodies were used to evaluate the molecular profile of interneurons, including glutamic acid decarboxylase (GAD), isoform 67 (localized in the neuronal cytoplasm [(Graus et al., 2020), see Fig. 3], involved in the synthesis of the inhibitory neurotransmitter GABA, calretinin (CR), and neuropeptide Y (NPY).

**Fig. 3.**
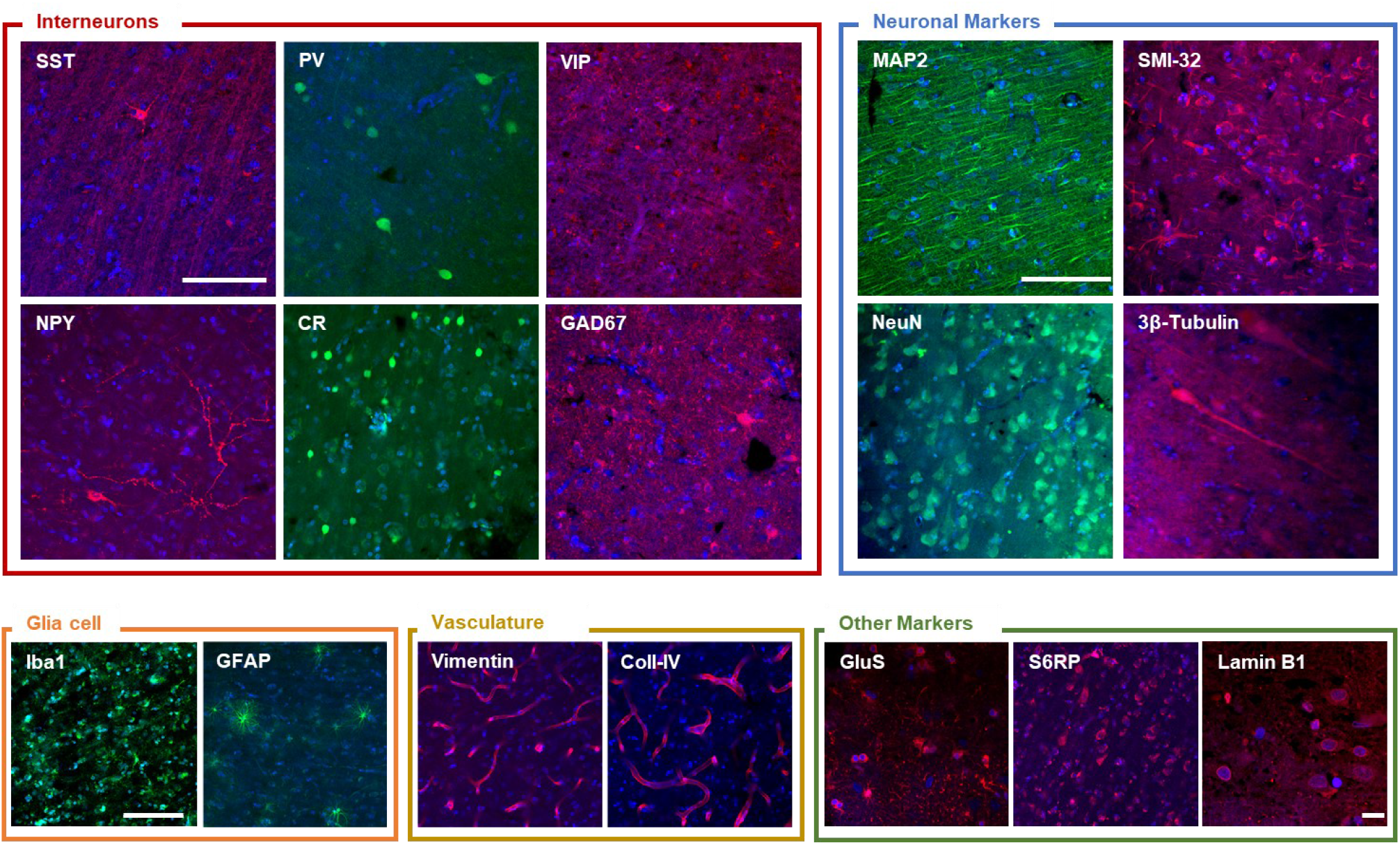
Antibody validation using confocal microscopy. Representative confocal images of the different validated staining. Interneurons: somatostatin (SST), parvalbumin (PV), vasointestinal peptide (VIP), neuropeptide Y (NPY), calretinin (CR), glutamic acid decarboxylase (GAD67). Neuronal markers: microtubule-associated protein 2 (MAP2), Non-phosphorylated neurofilament protein (SMI-32 antibody), neuron-specific nuclear protein (NeuN), 3β-Tub (3β-tubulin). Glial cell: ionized calcium binding adaptor molecule 1 (Iba1), glial fibrillary acidic protein (GFAP). Vasculature: vimentin, collagen IV (Coll-IV). Other markers: glutamine synthetase (GluS), phospho-S6 ribosomal protein (S6RP), lamin B1. Scale bar = 100 μm. Scale bar for lamin B1 image: 10 μm.

This assessment of interneurons populations confirmed that SWITCH/TDE is compatible with other markers that could be useful for the characterization of human brain tissue. We probed the microtubule-associated protein-2 (MAP2), the neurofilament protein, the neuronal nuclear antigen (NeuN), and β-tubulin. We extend the capability of this method with glial fibrillary acidic protein (GFAP) and ionized calcium-binding adapter molecule 1 (IBA1) (Fig. 3B). Also, phospho-ribosomal S6 ribosomal protein (p-S6RP), glutamine synthetase (GluS), and antibodies against vasculature (vimentin and collagen IV), and nuclear (lamin B1) components are compatible with this procedure (Fig. 3B), extending the possibility to investigate cytosolic, neurites, and nuclear markers in the human cortex.

## 4. Discussion

The SWITCH/TDE protocol allows clearing and labeling of thick adult human brain samples even after prolonged storage in formalin, making this method suitable to tissue stored in brain banks rather than to fresh or frozen tissue. Additionally, by exploiting the inter- and intramolecular bridges mediated by glutaraldehyde, SWITCH/TDE enables uniform preservation of the tissue architecture and molecular antigenicity. Here we demonstrate that SWITCH/TDE coupled with confocal microscopy can be an excellent method for probing different antibodies in human brain specimens and understanding its structural and cellular characteristics, making the analysis (such as segmentation, connectomics) of such materials more accurate. Indeed, the confocal microscope, nowadays a widely used instrument, with its optical sectioning capability is a key tool for antibody validation and 3D neuronal characterization.

However, several constraints reduce the power of the labeling and the quality of the imaging in human brain samples. These limitations are caused by glutaraldehyde used during the clearing process, as well as by the inherent characteristics of human brain samples. The glutaraldehyde used in the early stages of the SWITCH/TDE process is a strong crosslinker, characterized by two aldehyde groups connected by three carbons, which reacts with free amino groups located at the end of a polypeptide and side chain of lysines (Hopwood, 1972). This efficient ability to generate intra- and intermolecular bridges, fundamental for tissue preservation, can compromise epitopes, resulting in the loss of antibody immunoreactivity.

Moreover, natural autofluorescence of human brain samples due to lipopigments, cofactors, and blood vessels as well as the increase in autofluorescence caused by the prolonged fixation step of post-mortem samples create spurious signals that need to be considered. For instance, LF is the most abundant autofluorescence pigment in the human brain, localized in the perinuclear cytoplasm, and occasionally in the dendrites, axons, and presynaptic areas (Double et al., 2008). It is referred to as aggregates of undigested materials, characterized by the accumulation of oxidized proteins, lipids, and carotenoids, which affect neuron viability (Moreno-García et al., 2018). The presence of this autofluorescent pigment often leads to errors in the analysis of fluorescence images. Although several protocols have been developed for masking the autofluorescence signal of different lipopigments such as UV bleaching (Neumann and Gabel, 2002), Sudan Black (Oliveira et al., 2010), and sodium borohydride (Murray et al., 2015), they are potentially damaging to the tissue and time-consuming, leading to loss of antigens and, hence, immunolabeling sensitivity. For these reasons finding a proper way to discriminate such lipopigments from immunolabeling is essential. In this work, we assessed the autofluorescence distribution in the neuronal body by an easy method to distinguish the signal produced by LF with respect to specific immunostaining. Characterizing the emission spectrum of such lipopigments is an advantageous method for optimizing multiple staining at different excitation wavelengths.

Another challenge is to evaluate the antibody compatibility for both neuronal and non-neuronal markers in human brain samples. This is attributable to the high variability of post-mortem fixation conditions and epitope alterations in such samples. We present a list of validated antibodies that were screened using confocal microscopy on different human brain samples resulting in reliable staining. Such antibodies could be used for investigating a broad spectrum of biological questions in the human cortex. For example, in the adult central nervous system, class III beta-tubulin is typically neuron-specific, although altered expression patterns are noted in brain tumors. In gliomas, its expression is associated with an ascending grade of histologic malignancy (Katsetos et al., 2003). Instead, NeuN labeling shows high specificity for all neurons, and it is considered a late maturation marker during neuronal differentiation (Sarnat, 2013).

The characterization of glial cells allows obtaining information about inflammatory processes. For example, Hovens et al. demonstrated that in rats, IBA-1 could be used to evaluate microglial activation by analyzing cell body to cell size ratio (Hovens et al., 2014). Vasculature (vimentin and collagen IV) and nuclear (lamin B1) markers permit to study alterations of the anatomical organization of the tissue, for example after a stroke, or investigating chromatin organization (Bianchini et al., 2021) and neuronal migration during brain development (Lee et al., 2014). Finally, S6RP is a downstream effector of mTOR and its level of phosphorylation reflects the activation of the mTORC1 complex. It was demonstrated that S6RP is a potential disease-related gene in hemimegalencephaly with intractable epilepsy (Pelorosso et al., 2019). GluS is an ATP-dependent enzyme that catalyzes the condensation of glutamate with ammonia to yield glutamine. In the brain, GluS is primarily found in astrocytes, although changes in its activity or gene expression have been found in various neurological conditions (Jayakumar and Norenberg, 2016). It should be noted, however, that for many of these cellular markers, the penetration depth when using confocal is limited, and the whole sample is exposed to the excitation light, causing fluorophore bleaching during the acquisition. For these reasons, for 3D mesoscopic reconstruction, other optical techniques, like two-photon microscopy (Costantini and Mazzamuto et al., 2021) or light-sheet fluorescence microscopy (Gavryusev et al., 2019), are the best candidates for imaging such samples.

Nevertheless, we believe that the characterizations performed in this study can be exploited in future studies of the fine structural anatomy of the human brain, both in normal and pathological conditions. The demonstrated compatibility of the SWITCH/TDE method with different molecular markers allows both cell census and connectome analysis in the human brain specimens, permitting the broad reconstruction of neuronal spatial distribution patterns and discriminating the different populations of cells in the human brain.

## Acknowledgments

We thank Leah Morgan and Bruce Fischl, Massachusetts General Hospital, A.A. Martinos Center for Biomedical Imaging, Department of Radiology, USA for providing the human brain specimen 2 analyzed in this study. We express our gratitude to the donor involved in the body donation program of the Association des dons du corps du Centre Ouest, Tours, and of the Massachusetts General Hospital who made this study possible by generously donating his body to science. This project has received funding from the European Union’s Horizon 2020 research and innovation Framework Programme under grant agreement No. 654148 (Laserlab-Europe), from the European Union’s Horizon 2020 Framework Programme for Research and Innovation under the Specific Grant Agreement No. 785907 (Human Brain Project SGA2) and No. 945539 (Human Brain Project SGA3), from the General Hospital Corporation Center of the National Institutes of Health under award number U01 MH117023, and from the Italian Ministry for Education in the framework of Euro-Bioimaging Italian Node (ESFRI research infrastructure). Finally, this research was carried out with the contribution from “Fondazione CR Firenze” (private foundation). The content of this work is solely the responsibility of the authors and does not necessarily represent the official views of the National Institutes of Health.

